# Episodic memory retrieval affects the onset and dynamics of evidence accumulation during value-based decisions

**DOI:** 10.1101/2022.04.26.489332

**Authors:** Peter M. Kraemer, Sebastian Gluth

## Abstract

In neuroeconomics, there is much interest in understanding simple value-based choices where agents choose between visually presented goods, comparable to a shopping scenario in a supermarket. However, many everyday decisions are made in the physical absence of the considered goods, requiring agents to recall information about the goods from memory. Here, we asked whether and how this reliance on an internal memory representation affects the temporal dynamics of decision making on a cognitive and neural level. Participants performed a remember-and-decide task, in which they made simple purchasing decisions between money offers and snack items while undergoing EEG. Snack identity was presented either visually (value trials) or had to be recalled from memory (memory trials). Behavioral data indicated comparable choice consistency across both trial types, but considerably longer response times (RT) in memory trials. Drift-diffusion modeling suggested that this RT difference was due to longer non-decision time of decision processes as well as altered evidence accumulation dynamics (lower accumulation rate and higher decision threshold). The non-decision time effect was supported by a delayed onset of the lateralized readiness potential. These results show that both, decision and non-decision processes are prolonged when participants need to resort to internal memory representations during value-based decisions.

## 1 Introduction

Since its emergence as a scientific discipline more than two decades ago, neuroeconomics has often used simple consumer choices to investigate the cognitive and neural basis of value-based decision making (Glimcher et al., 2009). These choices are usually studied in a supermarket scenario, in which participants are exposed to visually presented choice sets of consumer goods and choose among them (e.g., Webb et al., 2021; Krajbich et al., 2010; Polanía et al., 2015; Frömer et al., 2019; Bakkour et al., 2019). While this scenario offers a well-controlled experimental setting, it has the limitation of only considering choices with the options being visually present. However, many value-based choices – for instance, when writing a shopping list or considering where to have lunch – are made in the physical absence of choice options (Lynch and Srull, 1982). In these cases, choice sets need to be created from memory and the cognitive and neural decision processes need to draw on internal memory representations (Stewart et al., 2006; Zhao et al., 2021). Thus, the aim of our study was to examine how this reliance on internal memory representations affects value-based decision processes.

Value-based decision making is often conceptualized with a two-stage model of valuation and action selection (e.g., Platt and Plassmann, 2013; Kable and Glimcher, 2009). In brief, this model distinguishes between valuation processes, where subjective value of choice options is integrated, and action selection, where different action plans compete until the winning plan is executed. Meta-analytic evidence from brain imaging studies demonstrated that valuation is strongly associated with activity in the ventromedial prefrontal cortex (vmPFC) (Bartra et al., 2013; Clithero and Rangel, 2014). This region seems to encode subjective value of choice options in various choice tasks, including the supermarket scenario. When decisions are based on memory representations, valuation seems to depend on the co-activation of the vmPFC and brain regions that account for memory retrieval, such as the hippocampus (Gluth et al., 2015) and the anterior prefrontal cortex (Zhang et al., 2021). While these studies focus on the effect of memory on valuation, the effect of memory representation on action selection processes have rarely been studied.

Action selection is frequently described as a competitive activation process in fronto-parietal brain regions in which the activity level of neural populations represent different action plans such as, approaching one vs. another choice option (Cisek, 2007; Gold and Shadlen, 2007). Over time, the activity levels change as a function of incoming valuation signals (Hare et al., 2011; Grueschow et al., 2015; Gluth et al., 2012). As soon as the activity level of one action plan outweighs the others by some critical margin, the according action is triggered and executed by the motor system (Thura and Cisek, 2014). The dynamic and competitive nature of action selection can be described by cognitive models of the sequential sampling framework (Shadlen and Kiani, 2013; Smith and Ratcliff, 2004). These models describe decision making as a noisy accumulation of *evidence* (see Fig. 1C). Here, evidence is an abstract unit of preference for either choice option. Over time, relative evidence accumulates until a threshold level is reached which – analogously to the neural process of action selection – triggers a corresponding action. Thereby the time to make a decision, the response time (RT), depends on the duration of the evidence accumulation process as well as on the non-decision time which is thought to include the duration of pre-decisional processes such as perceptual encoding and memory retrieval, as well as post-decisional processes such as motor execution (Ratcliff and McKoon, 2008; Ratcliff et al., 2016). Applied to economic choices such as consumer choices, sequential sampling models have received wide acceptance in neuroeconomics and decision neuroscience (Clithero, 2018; Fehr and Rangel, 2011; Busemeyer et al., 2019).

**Figure 1:**
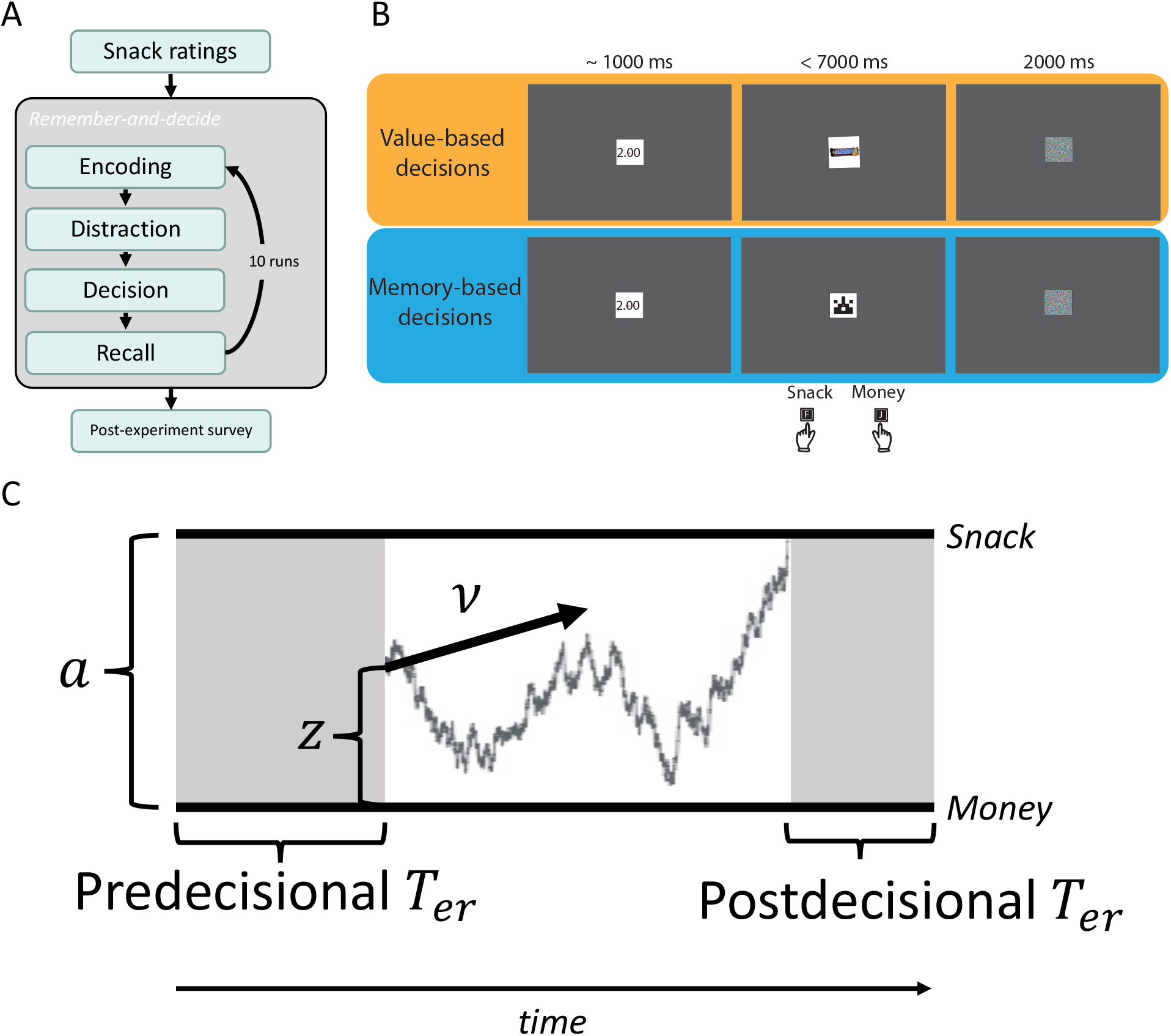
**A** Flow-chart of the experimental periods of the remember-and-decide task. **B** Illustration of the stimulus sequence of a value trial (orange) and a memory trial (blue). In each trial, participants chose between a monetary value and a snack option, which was presented either directly (value trial) or indicated by a unique identicon (memory trial). **C** Illustration of the dynamic choice process. Evidence accumulates over time with a drift-rate *υ*. The separation of the decision boundaries is defined by the parameter *a*. Here, the boundaries represent the two choice options of the remember-and-decide task (snack vs. money). The a-priori starting point bias is represented by the parameter *z*. Pre- and post-decisional non-decision time *T*_*er*_ are depicted in grey.

We investigated the neural dynamics of action selection by leveraging the temporal precision of electroencephalography (EEG). Specifically, we focused on the lateralized readiness potential (LRP) as a neural marker for action selection. The LRP is an effector-specific EEG component, recorded over the left and right motor cortices, which describes a rising negativity over several hundreds of milliseconds, reaching its peak shortly before an action is executed (Coles and Gratton, 1988). Although the LRP has traditionally been viewed as a pure motor component, there is good evidence that its propagation can last for more than a second, and that it can track the onset and the progression of decision processes from the presentation of choice-relevant information until a choice is made Gluth et al. (2013); Polanía et al. (2014). Furthermore, response-locked LRP analyses showed that the component follows a time course which resembles an accumulation-to-bound process of sequential sampling models (Schurger et al., 2012; Polanía et al., 2014). However, a detailed relation between LRP properties and evidence accumulation parameters is yet to be established. Here, we hypothesized that the onset, the slope and the peak amplitude of the LRP may be related to the onset, rate and threshold level of evidence accumulation, respectively.

We utilized this approach to compare the cognitive and neural dynamics of consumer choices, where goods were either presented visually vs. were recalled from memory. Cognitive modeling revealed major differences in the dynamics between both kinds of choices: Memory-based decisions yielded a lower rate of evidence accumulation, a higher decision threshold level as well as a delayed decision onset. Ultimately, the integration of different LRP properties into a neurally-informed sequential sampling model allowed us to identify the computational mechanisms of action selection in memory-based decisions.

## 2 Results

To study the differences of value-based and memory-based decisions, we invited hungry participants to take part in a laboratory study. In a remember-and-decide task, participants (N=39) cycled through several experimental periods (see Fig. 1B). During encoding, they associated abstract symbols (identicons) with individual snacks. After a 2-back distraction task, they made incentivized choices between money and snack offers (see Fig. 1C). Some trials (henceforth “value trials”) mimicked the supermarket scenario, with snacks being visually accessible. In other trials (henceforth “memory trials”), participants saw identicons and had to retrieve the associated snack identities from memory. Thus, in memory trials, their decisions relied on an internal memory representation of the snacks.

### 2.1 Behavioral results

Figure 2 depicts the average choice and RT data of the remember-and-decide task. For statistical analysis, we fitted hierarchical Bayesian logistic and linear regression models to the choice data (choice model) and RT data (RT model), respectively. While the choice model provided strong evidence that memory and value trials did not differ with respect to choice behavior (*β*_*Memory*_: 95% HDI = [−0.134, 0.096], *BF*_10_ = 0.06, Fig. 2B), the RT model provided decisive evidence that memory-based choices took longer (on average 395 ms; *sd* = 213 ms) than value-based choices (*β*_*Memory*_: 95% HDI = [0.265, 0.313], *BF*_10_ = 1.34 * 10^123^, Fig. 2D).

**Figure 2:**
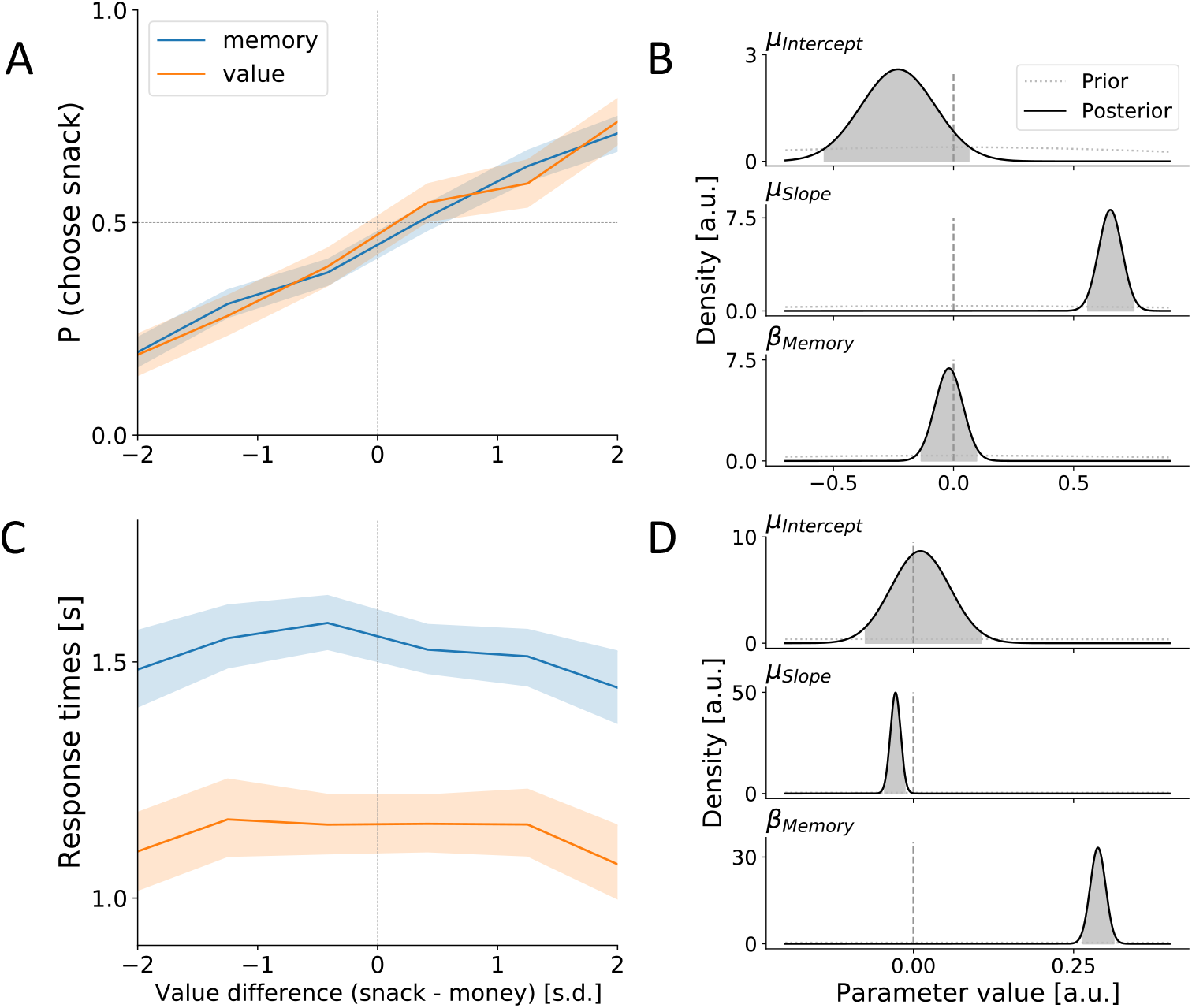
Behavioral results. **A** Mean proportion of snack choices as a function of value difference. Shaded areas indicate the bootstrapped 95% confidence intervals. **B** Logistic regression estimates of choices. Full lines indicate the posterior probability distribution of relevant parameters (*µ*_*Intercept*_ and *µ*_*Slope*_ for intercept and slope of the regression line; *β*_*Memory*_ models the difference between memory trails and value trials). Shaded areas indicate the 95% highest density interval. Dotted lines indicate the prior probability distribution. Vertical dashed lines indicate the parameter value specifying the null hypothesis. **C** Mean response times as a function of value difference. **D** Linear regression estimates of response times.

In addition to the effect on RT, we found decisive evidence that choice behavior depended on value difference of the choice options (logistic model, *µ*_*Slope*_: 95% HDI = [0.559, 0.751], *BF*_10_ = 1.80 * 10^37^, Fig. 2B), and substantial evidence that RT decreased as a function of absolute value difference (linear model, *µ*_*Slope*_: 95% HDI = [−0.045, −0.013], *BF*_10_ = 3.66, Fig. 2D). Both results are in line with core predictions of sequential sampling models (Clithero, 2018).

We found strong evidence that RT did not differ depending on whether the snack or the money was chosen (linear model, *µ*_*Intercept*_: 95% HDI = [−0.075, 0.106], *BF*_10_ = 0.05, Fig. 2D), and anecdotal evidence for unbiased choice behavior (logistic model, *µ*_*Intercept*_: 95% HDI = [−0.538, 0.064], *BF*_10_ = 0.47, Fig. 2B), both suggesting the absence of an a-priori preference for either option (hence, a starting-point bias in DDM terminology; cf. Lopez-Persem et al., 2016).

### 2.2 Cognitive modeling

The central findings of our behavioral analyses were that participants had similar choice consistency in value and memory trials but took significantly longer to execute memory-based choices. Given that choice consistency and response times are inherently related in the sequential sampling framework, this pattern of results can be explained by two distinct, not mutually exclusive mechanisms: First, increased RT may result from increased non-decision time and a corresponding shift in RT while leaving choice consistency unaffected. A second explanation may be prolonged decision time, caused by altered evidence accumulation dynamics. For instance, increased boundary separation has been linked to increased RT while affecting choice coherence only moderately (Fontanesi et al., 2019).

We investigated these potential explanations by fitting a drift-diffusion model (DDM) to the behavioral data. The model is depicted in Figure 3 and fully specified in the Methods (section 4.7). Importantly, while individual variance of the diffusion parameters is explained by random effects, the difference between value trials and memory trials is modeled with fixed effects *δ* parameters.

**Figure 3:**
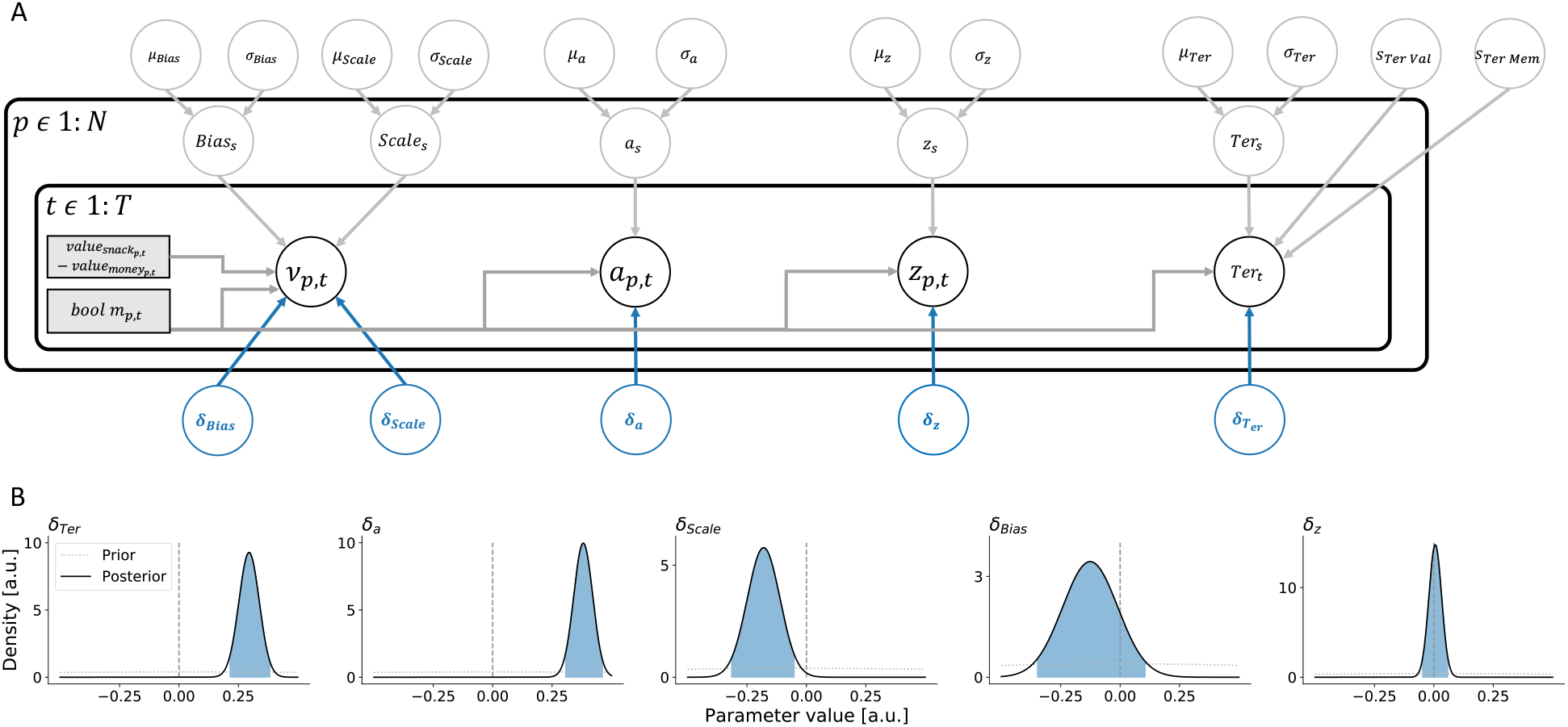
Cognitive modeling. **A** Graphic illustration of the structure of the drift-diffusion model. Parameters are depicted in circles, variables in rectangles. Arrows indicate the inter-relation of parameters and variables. Indices represent the participant level p and the trial level t. **B** Probability distributions for *δ* parameters (with positive values indicating higher parameter estimates for memory compared to value trials). Full lines indicate the posterior probability distribution of relevant parameters. Shaded areas indicate the 95%-highest density interval. Dotted lines indicate the prior probability distribution. Vertical dashed lines indicate the parameter value which specifies the null hypothesis.

We found decisive evidence that participants exhibited a longer non-decision time during memory trials (*δ*_*Ter*_ : 95%-HDI: [0.22, 0.38], *BF*_10_ = 7.14 * 10^8^). Concurrently, the dynamics of the diffusion process seemed to be altered. The boundary separation in memory trials was estimated credibly larger than in the value condition (*δ*_*a*_ : 95%-HDI: [0.31, 0.46], *BF*_10_ = 2.01 * 10^18^). Additionally, the rate of evidence accumulation, as governed by the *Scale* parameter, may be lower in memory trials, although evidence is relatively weak: While the 95%-HDI ([-0.32,-0.05]) excluded zero, the Bayes factor of 2.07 only provided anecdotal evidence for this lower drift scale.

With respect to potential biases in the diffusion process, we found that neither the starting point bias *µ*_*z*_, nor a drift-rate bias *µ*_*Bias*_ (which models a general preference for snacks or money during the accumulation process Kraemer et al., 2021a; Krajbich, 2021) were credibly different from zero as indicated by the 95% −HDIs of these parameters (see Table 1). Furthermore, we found moderate and very strong evidence against potential biases in memory trials as compared to value trials (*δ*_*z*_ : 95%-HDI: [−0.05, 0.06], *BF*_10_ = 0.03; *δ*_*Bias*_ : 95%-HDI: [−0.35, 0.11], *BF*_10_ = 0.21), letting us conclude that diffusion processes had no general preference for money or snack choices.

**Table 1:** Parameter estimates of the cognitive drift-diffusion model.

| | Prior | Posterior M | Posterior sd | 95%-HDI | $BF_{10}$ |
| --- | --- | --- | --- | --- | --- |
| $\mu_{Ter}$ | $\mathcal{N}(-1, 1)$ | -0.754 | 0.120 | [-0.99,-0.52] | - |
| $\sigma_{Ter}$ | $\mathcal{HN}(0, 1)$ | 0.707 | 0.091 | [0.54,0.89] | - |
| $\delta_{Ter}$ | $\mathcal{N}(0, 1)$ | 0.295 | 0.043 | [0.22,0.38] | $7.14 * 10^8$ |
| $s_{Ter, Value}$ | $\mathcal{HN}(0, 1)$ | 0.392 | 0.028 | [0.34,0.44] | - |
| $s_{Ter, Memory}$ | $\mathcal{HN}(0, 1)$ | 0.446 | 0.021 | [0.4,0.49] | - |
| $\mu_a$ | $\mathcal{N}(2, 1)$ | 1.521 | 0.081 | [1.37,1.68] | - |
| $\sigma_a$ | $\mathcal{HN}(0, 3)$ | 0.457 | 0.058 | [0.35,0.57] | - |
| $\delta_a$ | $\mathcal{N}(0, 1)$ | 0.381 | 0.040 | [0.31,0.46] | $2.01 * 10^{18}$ |
| $\mu_{Scale}$ | $\mathcal{N}(0, 1)$ | -0.923 | 0.089 | [-1.1,-0.76] | - |
| $\sigma_{Scale}$ | $\mathcal{HN}(0, 1)$ | 0.406 | 0.068 | [0.28,0.54] | - |
| $\delta_{Scale}$ | $\mathcal{N}(0, 1)$ | -0.180 | 0.069 | [-0.32,-0.05] | 2.07 |
| $\mu_{Bias}$ | $\mathcal{N}(0, 1)$ | -0.291 | 0.211 | [-0.69,0.14] | - |
| $\sigma_{Bias}$ | $\mathcal{HN}(0, 1)$ | 1.174 | 0.160 | [0.88,1.49] | - |
| $\delta_{Bias}$ | $\mathcal{N}(0, 1)$ | -0.126 | 0.116 | [-0.35,0.11] | 0.21 |
| $\mu_z$ | $\mathcal{N}(0, 1)$ | 0.026 | 0.035 | [-0.04,0.09] | - |
| $\sigma_z$ | $\mathcal{HN}(0, 1)$ | 0.161 | 0.026 | [0.11,0.21] | - |
| $\delta_z$ | $\mathcal{N}(0, 1)$ | 0.006 | 0.027 | [-0.05,0.06] | 0.03 |

The table lists the respective prior distributions, mean of the posterior distribution, standard deviation of the posterior distribution, the 95%-Highest density interval, and, where applicable, the Bayes factor.

### 2.3 EEG results

The cognitive modeling analyses suggested altered diffusion dynamics (boundary separation and drift scale) as well as an increased non-decision time in memory trials. We next asked whether these findings are supported by an analysis of the simultaneously recorded LRP. We focused on three LRP properties: The stimulus-locked LRP onset as a marker for the end of predecisional non-decision time, the response-locked LRP slope as a marker for speed of evidence accumulation, and the response-locked peak amplitude as a marker for decision threshold.

Figure 4A shows the stimulus-locked LRP waveforms for both conditions. We estimated the LRP onset time for both conditions using a hierarchical Bayesian segmented-regression method. Figure 4C depicts the difference of the LRP onset between memory and value trials as the difference in their posterior probability estimates. Calculating the proportion of posterior samples larger than 0 indicated that there is probability of 98.1% that memory trials had a later LRP onset than value trials, given the model and the data. The unit of parameter values in Figure 4C can also be interpreted as units of seconds. Thereby, the average difference in non-decision time amounts to 83 ms (*sd* = 41ms). This finding suggests that the delayed LRP onset may account for a part of the 395 ms RT difference between the conditions.

**Figure 4:**
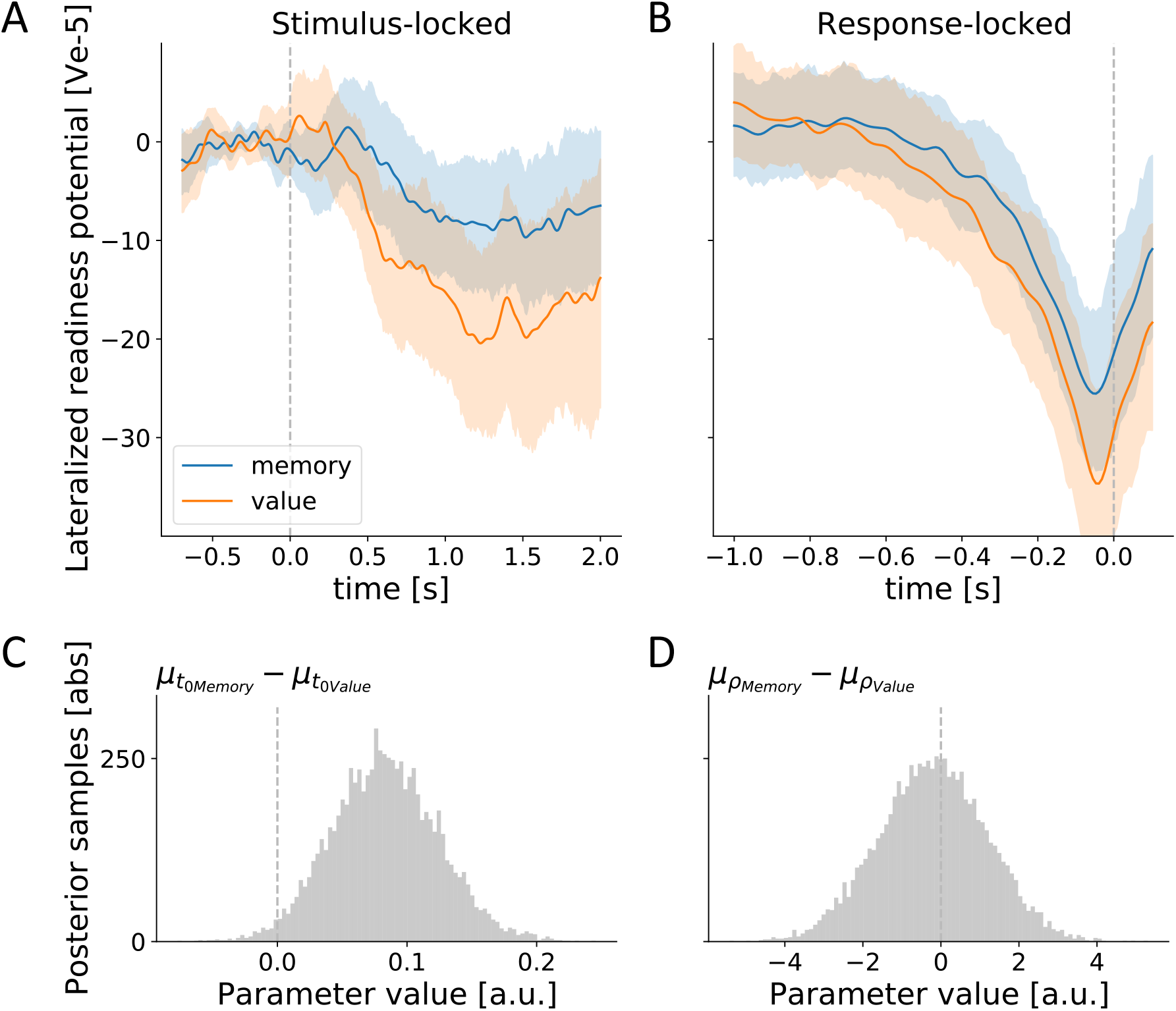
LRP analyses. **A** Mean stimulus-locked LRP waveforms for both conditions. Shaded areas indicate 95% bootstrapped confidence intervals. **B** Mean response-locked LRP waveforms. **C** Posterior difference in LRP onset between memory and value trials. Histogram shows posterior samples from group-level onset estimates of a hierarchical Bayesian segmented-regression model. Vertical dashed line indicate onset equivalence. **D** Posterior difference in response-locked LRP slope between memory and value trials.

Analysis of the response-locked LRP properties did not indicate differences between value and memory trials. The ratio of posterior difference samples for LRP slope only indicated a 44.3% probability for a larger slope in memory trials. A Bayesian t-test of LRP peak amplitudes yielded substantial evidence against a difference between conditions (*t*(38) = 0.68; *BF*_10_ = 0.22).

In sum, we found credible evidence that memory trials have a later LRP onset, which partially explains the behavioral difference in RT. The properties which we hypothesized to relate to evidence accumulation (i.e. slope and peak amplitude), however, were not different between conditions.

### 2.4 Neuro-cognitive modeling

In a next step, we sought to relate the cognitive modeling and EEG results by testing whether the different LRP properties can inform the parameters of the cognitive model. To answer this question, we re-estimated our DDM and added fixed effects *θ* parameters which allowed us to test, whether the individual variability of LRP properties explained variance of the diffusion parameters on the participant level. The structure of the neuro-cognitive DDM is displayed in Figure 5. As indicated in green, the *θ* parameters were allowed to affect the scale parameter of the drift-rate (*θ*_*Slope*_), the boundary separation (*θ*_*Peak*_) and the non-decision time (*θ*_*Onset*_) on the participant level (see 4.8 for details).

**Figure 5:**
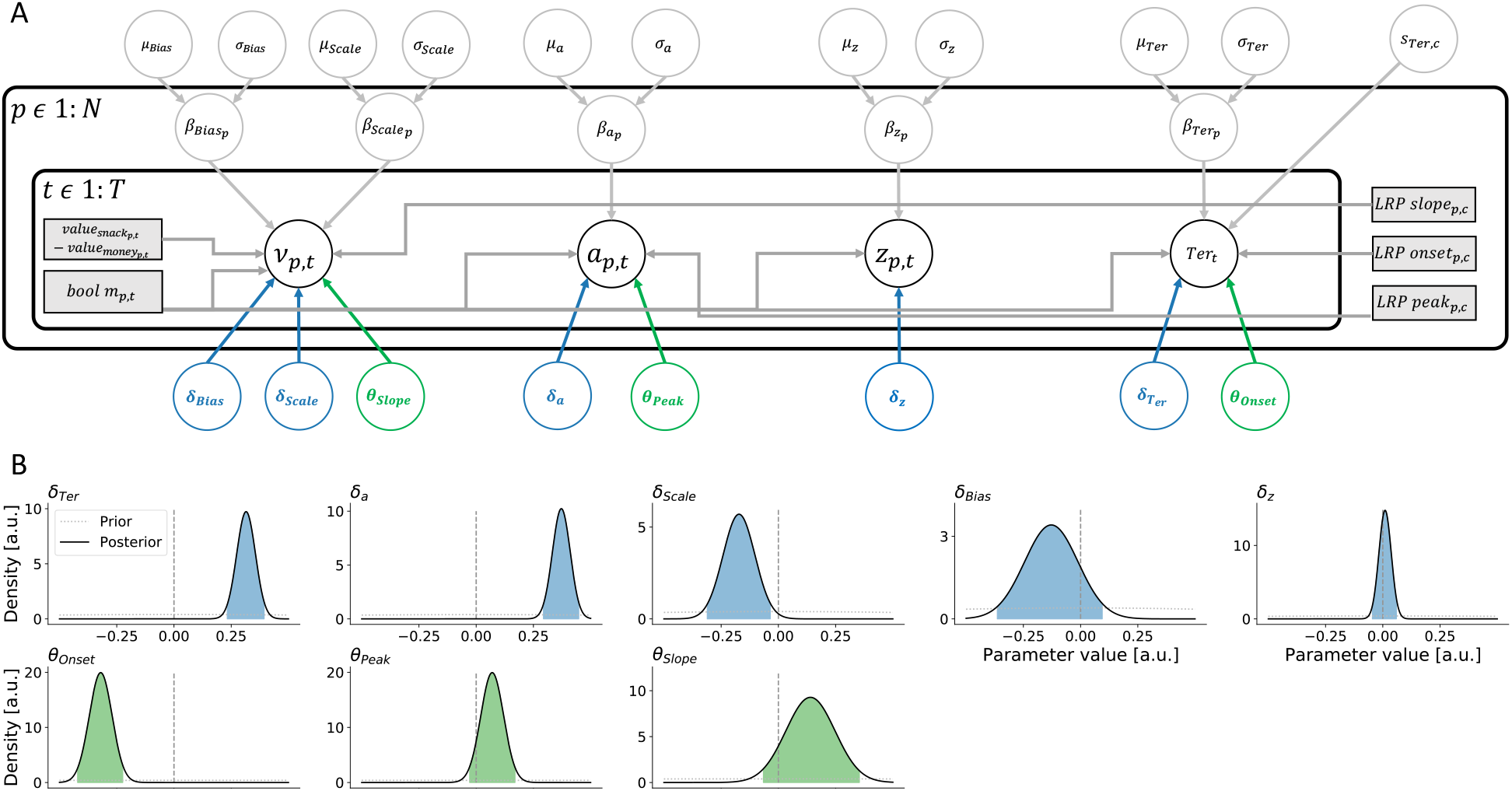
Neuro-cognitive modeling. **A** Graphic illustration of the structure of the drift-diffusion model. Parameters are depicted in circles, variables in rectangles. Arrows indicate how parameters and variables are related. Indices represent the participant level p and the trial level t. **B** Probability distributions for *δ* and *θ* parameters. Full lines indicate the posterior probability distribution of relevant parameters. Shaded areas indicate the 95%-highest density interval. Dotted lines indicate the prior probability distribution. Vertical dashed lines indicate the parameter value which specifies the null hypothesis.

Prior and posterior parameter distributions can be inspected in Table 2. Consistent with the purely cognitive DDM, the results of estimating the neuro-cognitive DDM suggest that memory trials were associated with larger non-decision time (*δ*_*Ter*_ : 95%-HDI: [0.23, 0.39], *BF*_10_ = 2.69 * 10^11^), larger boundary separation (*δ*_*a*_ : 95%-HDI: [0.29, 0.45], *BF*_10_ = 2.22 * 10^18^) and (moderate evidence for) decreased rate of evidence accumulation (*δ*_*Scale*_ : 95%-HDI: [−0.31, −0.04], *BF*_10_ = 1.43). Again, diffusion processes were unbiased between snack and money choices (*µ*_*z*_ : 95%-HDI: [−0.04, 0.1]; *µ*_*Bias*_ : 95%-HDI: [−0.69, 0.14]).

**Table 2:**
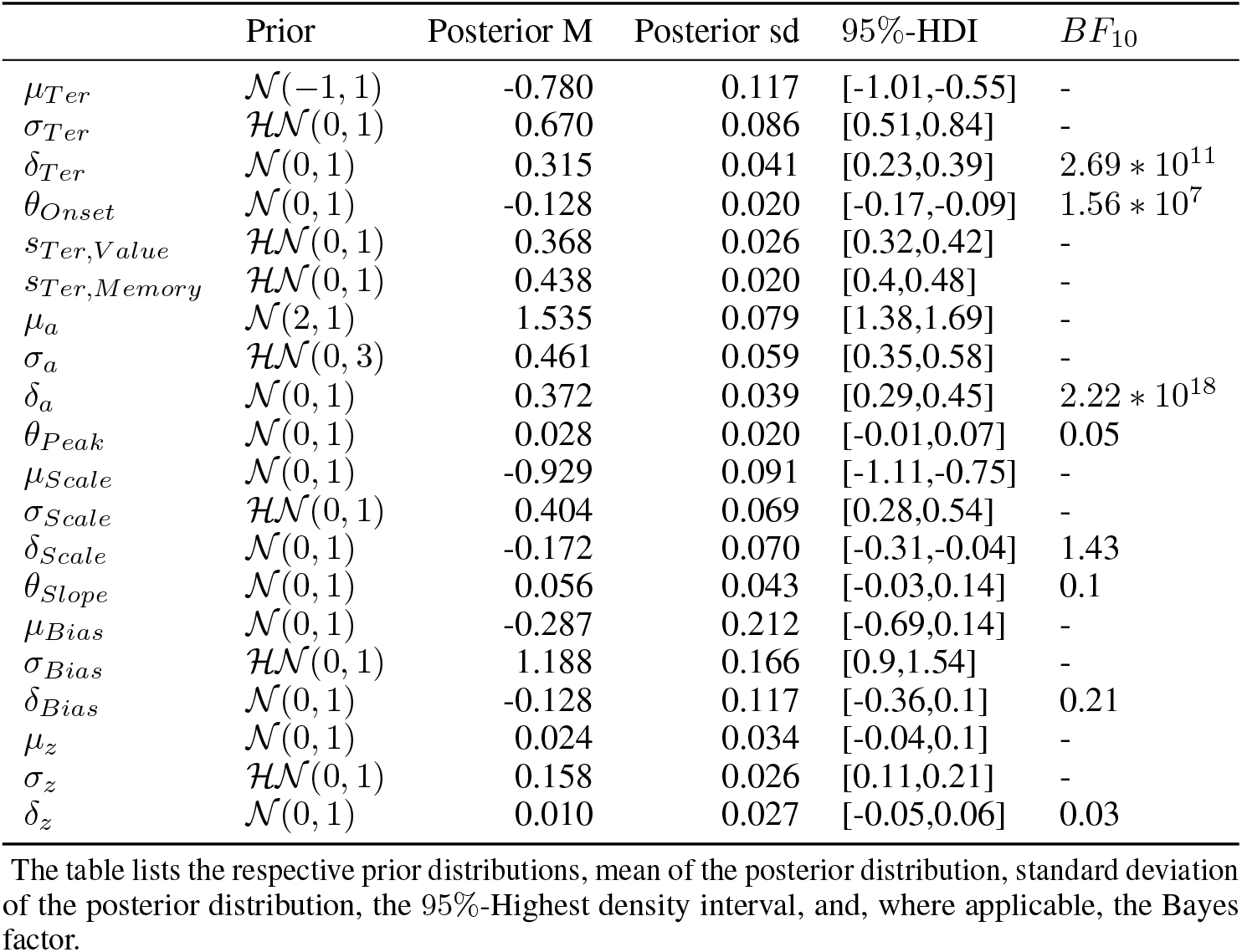
Parameter estimates of the neuro-cognitive drift-diffusion model.

| | Prior | Posterior M | Posterior sd | 95%-HDI | $BF_{10}$ |
| --- | --- | --- | --- | --- | --- |
| $\mu_{Ter}$ | $\mathcal{N}(-1, 1)$ | -0.780 | 0.117 | [-1.01,-0.55] | - |
| $\sigma_{Ter}$ | $\mathcal{HN}(0, 1)$ | 0.670 | 0.086 | [0.51,0.84] | - |
| $\delta_{Ter}$ | $\mathcal{N}(0, 1)$ | 0.315 | 0.041 | [0.23,0.39] | $2.69 * 10^{11}$ |
| $\theta_{Onset}$ | $\mathcal{N}(0, 1)$ | -0.128 | 0.020 | [-0.17,-0.09] | $1.56 * 10^7$ |
| $s_{Ter,Value}$ | $\mathcal{HN}(0, 1)$ | 0.368 | 0.026 | [0.32,0.42] | - |
| $s_{Ter,Memory}$ | $\mathcal{HN}(0, 1)$ | 0.438 | 0.020 | [0.4,0.48] | - |
| $\mu_a$ | $\mathcal{N}(2, 1)$ | 1.535 | 0.079 | [1.38,1.69] | - |
| $\sigma_a$ | $\mathcal{HN}(0, 3)$ | 0.461 | 0.059 | [0.35,0.58] | - |
| $\delta_a$ | $\mathcal{N}(0, 1)$ | 0.372 | 0.039 | [0.29,0.45] | $2.22 * 10^{18}$ |
| $\theta_{Peak}$ | $\mathcal{N}(0, 1)$ | 0.028 | 0.020 | [-0.01,0.07] | 0.05 |
| $\mu_{Scale}$ | $\mathcal{N}(0, 1)$ | -0.929 | 0.091 | [-1.11,-0.75] | - |
| $\sigma_{Scale}$ | $\mathcal{HN}(0, 1)$ | 0.404 | 0.069 | [0.28,0.54] | - |
| $\delta_{Scale}$ | $\mathcal{N}(0, 1)$ | -0.172 | 0.070 | [-0.31,-0.04] | 1.43 |
| $\theta_{Slope}$ | $\mathcal{N}(0, 1)$ | 0.056 | 0.043 | [-0.03,0.14] | 0.1 |
| $\mu_{Bias}$ | $\mathcal{N}(0, 1)$ | -0.287 | 0.212 | [-0.69,0.14] | - |
| $\sigma_{Bias}$ | $\mathcal{HN}(0, 1)$ | 1.188 | 0.166 | [0.9,1.54] | - |
| $\delta_{Bias}$ | $\mathcal{N}(0, 1)$ | -0.128 | 0.117 | [-0.36,0.1] | 0.21 |
| $\mu_z$ | $\mathcal{N}(0, 1)$ | 0.024 | 0.034 | [-0.04,0.1] | - |
| $\sigma_z$ | $\mathcal{HN}(0, 1)$ | 0.158 | 0.026 | [0.11,0.21] | - |
| $\delta_z$ | $\mathcal{N}(0, 1)$ | 0.010 | 0.027 | [-0.05,0.06] | 0.03 |

The table lists the respective prior distributions, mean of the posterior distribution, standard deviation of the posterior distribution, the 95%-Highest density interval, and, where applicable, the Bayes factor.

Most importantly, we found a positive relationship of LRP onsets and non-decision time which implies that participants who exhibit a later LRP onset also exhibit a larger non-decision time. This was indicated by *θ*_*Onset*_ being credibly smaller than zero (95%-HDI: [−0.17, −0.09], *BF*_10_ = 1.56 * 10^7^; parameter values are negative for purely mathematical reasons, see 4.8 for details). Our analysis showed that, not only does increased non-decision time in memory trials coincide with a delayed LRP onset on the population level, but also that the individual LRP onsets were related to individual estimates of non-decision time.

With respect to LRP peak amplitudes and LRP slope, we found evidence for the null hypotheses that they were unrelated to the boundary separation (*θ*_*Peak*_ : 95%-HDI: [−0.01, 0.07], *BF*_10_ = 0.05) and scale parameter (*θ*_*Slope*_ : 95%-HDI: [−0.03, 0.14], *BF*_10_ = .1), respectively. Thus, the altered diffusion dynamics (boundary separation and drift scale) could not be linked to individual LRP properties.

## 3 Discussion

We investigated the differences of action selection processes between value-based and memory-based decisions. Whereas participants exhibited choice behavior equally coherent with subjective value ratings, their RT were considerably longer in memory-based decisions. According to our diffusion model analysis, this difference was not attributed to a single but to multiple underlying mechanisms: on the one hand, we found a longer non-decision time, presumably related to memory retrieval processes, and on the other hand there was a longer accumulation-to-bound process as described by both a larger boundary separation and a (potentially) smaller drift-rate. On the neural level, the finding of a longer non-decision time was supported by a delayed LRP onset in general as well as in a positive correlation between non-decision time parameter and LRP onset across participants. Contrary to our hypotheses, the slope and peak amplitude of the LRP were unrelated to evidence accumulation dynamics.

Prominent theories about memory-based decision making state that decisions are made by retrieving and integrating information bits as samples over time (Johnson et al., 2007; Stewart et al., 2006; Zhao et al., 2021). This idea is taken up by sequential models of memory-based decision making (Shadlen and Shohamy, 2016; Kraemer et al., 2022). According to our cognitive modeling efforts, participants applied a higher decision threshold to memory-based decisions, requiring them to sample information for a longer time period. At the same time, the rate of evidence accumulation, as indicated by the drift-scale parameter, was potentially lower, which suggests that memory-based information was less directive for the eventual decision outcome. We argue that these altered evidence accumulation dynamics may be linked to the additional cognitive demands of episodic retrieval in memory trials (Weilbächer et al., 2021). More specifically, when internal memory representations guide momentary evidence, individual samples may be less informative for the valuation and action selection processes. The decision processes adapt to this situation by sampling for a longer time, which can lead to similar choice consistency but longer RT.

In our study, we find clear indications that the LRP can be viewed as a neural marker of the onset of decision processes. While the LRP remains being mainly viewed as a motor component, it has originally been proposed as a “window on the mind and brain” (Coles, 1989). Coles and Gratton (1988) suggested to use the LRP as a neural marker of early communication, such that “when lateralization is observed, following stimulus presentation, the subject presumably has access to a particular aspect of the stimulus information” (p. 87). The LRP onset can therefore be considered as a measure of when external information from visual stimuli vs. internal information from memory begin to affect the action selection process (Gluth et al., 2013). Our findings are in line with this assumption as the delayed LRP onset coincided with a larger non-decision time estimate for memory trials. The onset of the decision process may thus be delayed due to additional time affordances of memory retrieval processes (note, however, that memory retrieval was likely not completed at the time of the LRP onset, see below). Concurrently, individual LRP onsets were informative to estimate non-decision times on the participant level, coherent with earlier findings in perceptual decision making (Van Vugt et al., 2014). These findings support the assumption that the LRP onset may serve as a neural marker for the end of pre-decisional non-decision time, and hence, the beginning of the evidence accumulation process. A more systematic test of this hypothesis is needed, for instance via an experimental manipulation of pre-decisional non-decision time and (Nunez et al., 2019) and the simultaneous measurement of LRP onsets. Nonetheless, the LRP onset bears great potential for estimating the time at which particular stimulus information is accessible to the neural system and can affect evidence accumulation, as has been discussed in recent neuroeconomic literature (Maier et al., 2020; Sullivan et al., 2015).

Whereas the LRP onset was related to non-decision time, the other LRP properties we considered were unrelated to evidence accumulation: Slope and peak amplitude were not different between value and memory trials and were not informative for the estimation of drift scale and boundary separation on the individual level. Thus, we reject our initial hypotheses which stated that these properties may reflect evidence accumulation dynamics in this direct way. Unfortunately, we therefore lacked a neural marker of evidence accumulation to back up the differences in drift scale and boundary separation, as indicated by cognitive modeling. Another attempt, focused at the centro-parietal positivity – a component which has been linked to evidence accumulation in perceptual (O’Connell et al., 2012; Kelly and O’Connell, 2013), and value-based decision making (Pisauro et al., 2017) – was rejected as we did not observe the expected response-locked ramping activity of a putative CPP in our data (See Appendix 8). Given that also other authors struggled to establish a close relationship between CPP and evidence accumulation (Lui et al., 2021; Frömer et al., 2021), we notice that the field lacks the knowledge of EEG components which reflect evidence accumulation over a wider range of tasks.

The two-stage model of valuation and action selection suggests two processes that contribute to decision making. Importantly, the model remains agnostic about whether valuation must have been terminated before action selection begins (Platt and Plassmann, 2013). Under this assumption of strictly serial processes, participants in our task would have completed episodic memory retrieval before starting the action selection process, because a proper value representation requires a precise memory of the choice option in our task. To investigate this, we aimed to identify neural signatures of memory retrieval and decision making, and to observe these signals as they unfold over time. However, we could not find a marker for (lateralized) memory signals as signs of replay, as other researchers did (Waldhauser et al., 2016, 2012; Ede et al., 2019) (Appendix 9.2). Nonetheless, based on the timing of the LRP onset, we speculate that memory retrieval, valuation and action selection processes are unlikely to occur in a strictly serial manner: In cued recall (as required in our task) memory reinstatement occurs between 500 and 1,500 ms (see, Staresina and Wimber, 2019, for a review), and as the LRP onset was estimated at 235 ms in our task, we reason that action selection likely started before the critical time period for retrieval was reached. Hence, memory retrieval and action selection processes at least seem to run in parallel over some time. This is consistent with models following the parallel distributed processing doctrine (Hunt and Hayden, 2017; Yoo and Hayden, 2018) which argue for non-sequential and instead a largely parallel processing during value-based decision making.

Moreover, the LRP onset results have important implications for sequential sampling models, which often assume that the duration of memory retrieval is related to (pre-decisional) non-decision time (Ratcliff and McKoon, 2008; Ratcliff et al., 2016; Shepherdson et al., 2018). Our findings support the notion that the requirement to retrieve relevant information from memory affects both, decision dynamics and non-decision time. How can future sequential sampling models account for memory-based decisions? First, non-decision time may be equipped with additional parameters which model the effects of memory retrieval, as we did in our study with the *δ*_*Ter*_ and *s*_*Ter*_ parameters. Second, evidence accumulation dynamics should account for the fact that memory retrieval is not an instantaneous event but rather a process where information gets more vivid over time (presumably in theta rhythm, Kerrén et al., 2018; Staresina and Wimber, 2019). Such retrieval processes could be embedded in sequential sampling models where the state of memory retrieval at a time causes non-linear accumulation dynamics, as have been proposed for conflict tasks (White et al., 2017) and in judgments drawing on semantic memory (van Maanen et al., 2012; Kraemer et al., 2021b). The validation of such models will crucially depend on the identification of temporally precise neural markers of the onset, continuation and termination memory and decision processes.

## 4 Methods

### 4.1 Participants

In total, 48 participants took part in the study. 9 participants had to be excluded due to dietary incompatibilites of the snacks (n=3), acute headache (n=1), problems in understanding or performing the tasks (n=2) or technical problems with the EEG system (n=3). Our final data set comprised 39 healthy, right-handed human volunteers (28 female, mean age = 22.5, age range = [19, 29]). Participants were recruited via online recruitment systems of the University of Basel and were reimbursed either with a money amount of 90 CHF or with course credits. Participants were instructed to fast for at least four hours before the start of the experiment. Additionally, the tasks were incentivized (see 4.2). The study was approved by the ethics committee of the University of Basel, and all participants gave written informed consent to the procedures.

### 4.2 Procedure

The study was conducted in two separate sessions (mean time between sessions = 7.5 days, range = [7, 15]). Session 1 was a training session, where participants were familiarized with the task structure. This was done to minimize learning effects in session 2, where the relevant data for this study was recorded. In both sessions, participants first rated 55 snack items regarding their willingness to pay. Next, they performed 10 runs of a remember-and-decide task. In a post-experiment survey, participants rated their subjective familiarity with the snacks and were interviewed about potential problems and personal strategies of the task procedures.

#### 4.2.1 Setup

In session 1, we collected behavioral data. In the laboratory, participants were seated in front of a screen (size = 47.38 × 29.61 cm ≙ 24′′ screen diagonal, resolution = 1680 × 1050 pixels, refresh rate = 60 Hz). Their heads were fixed with a chin rest at 60 cm distance to the screen. Eye tracking data was collected with an SMI iViewX RED 500 system (SensoMotoric instruments, Berlin, Germany) with 500 Hz sampling rate and SMI BeGaze (version 3.4) as recording software. Note that the purpose of using an eye tracker was to control and restrict eye movements rather than to investigate any potential effects of attention.

In session 2, we collected behavioral and EEG data. The setup was identical to session 1 with the exception that the SMI eye tracker in the EEG laboratory had a sampling rate of 60 Hz. EEG data was recorded with a 64 electrode ActiveTwo system (BioSemi, Amsterdam, The Netherlands) at a sampling rate of 1024 Hz. The recording software was BioSemi ActiView. Scalp electrodes were placed corresponding to the 10-10 system using elastic caps. To record EOG, we placed two electrodes temporal to the eyes and two electrodes above and below the right eye. EMG activity at both hands were recorded using two electrodes at the opponens pollicis and the flexor digitorum.

All experiments were programmed in Python using the exPyriment library (Krause and Lindemann, 2014, version 0.6.3). The eye-tracker was coupled to the experiment using PyGaze (Dalmaijer et al., 2014, version 0.6.0). Eye-tracking recording software was iView X (version 3.1.0). EEG recording software was ActiView (version 7.04).

#### 4.2.2 Snack ratings

After participants gave written informed consent and were instructed about the experimental procedures, they were familiarized with the snack stimuli. Participants first saw every snack and its name once. They were allowed to take each snack in their hand and read the information about the ingredients provided by the package. Afterwards we assessed their willingness to pay for each snack. Thereto, the snacks were presented on a screen and participants moved a slider continuous scale between 0 and 5 CHF to rate them. The slider was initialized on a middle position and had to be moved at least once to commit a rating. Snack order was randomized. Every snack was rated twice. Following previous studies (e.g., Gluth et al., 2015; Krajbich et al., 2010), we used the average rating of each snack as a proxy for subjective snack value. The task was incentivized as one snack rating was chosen randomly and compared to a random number from a uniform distribution between 0 and 5. The participants would receive the snack if its rating was higher than the random number, or the random number as money if it was higher than the snack rating.

#### 4.2.3 Remember-and-decide task

The remember-and-decide task consisted of ten consecutive runs in which participants completed the four periods encoding, distraction, decision, and recall (in that order).

During encoding, participants learned associations between abstract symbols (identicons) and snacks. In each trial, participants fixated on a red fixation dot within a small central square (size: 84 × 84 pixels) in the center, flanked by two large lateral squares (size: 320 × 320 pixels), one in every hemifield. After a minimum fixation time (jittered interval drawn from a uniform distribution between 966 and 1033 ms, controlled via online eye tracking), an identicon appeared in the center square alongside with a snack image in either lateral square. Participants memorized the association of each identicon with its respective snack. They maintained fixation throughout the trial. Fixation control was ensured by simultaneous eye tracking. When the gaze of the participants left the central square, the trial would abort with a message calling out the fixation break, shown for 3 s. Participants were further informed that every fixation break would reduce their chance of winning a snack after the experiment by 2 %. After 2000 ms the squares were superseded by visual masks as inter-trial intervals in which participants were allowed to blink. In every run, participants learned 5 identicon-snack associations. Each identicon-snack association was presented several times. To investigate the effect of presentation times on learning performance, the associations were presented 3 or 5 times (counterbalanced across runs) in session 1 and 4 times in session 2. To avoid random clustering of specific identicon-snack associations at a specific time window in the encoding period (e.g., that the identicon-Snickers association is presented several times at the end of the encoding period), we pseudo-randomized their presentation order so that all associations were presented in mini-blocks (not noticeable to the participants) within which the order was fully randomized.

During distraction, participants performed a 2-back task so that they could not rehearse information obtained in the encoding phase. For 30 s, digits were presented in the center of the screen. Participants were instructed to press the space button on their keyboard when the current digit was identical to the penultimate digit. Digits were presented for 900 ms with an inter-trial interval of 100 ms. The order of digits was pseudo-randomized so that on average one third of the digits were equal to the penultimate digit.

During the decision period, participants made choices between snacks and money amounts. In each decision trial, participants first saw a money offer presented in a white square (size: 84 × 84 pixels) with a red fixation dot. Participants were required to look at the fixation dot throughout the whole trial. After a minimum fixation time (again, jittered interval as during encoding), the money stimulus was superseded by a snack stimulus. The snack stimulus could either be a snack image (value trials) or an identicon (memory trials), learned during the encoding period. Only in value trials, the snack identity was visually accessible. In memory trials, it had to be retrieved from memory in order to make an informed choice. Participants were free to choose either the money amount or the snack by pressing either f vs. j button with their left vs. right index finger. Button-option assignment was counter-balanced across participants. If a maximum decision time of > 7000 ms was reached, the trial aborted calling out that no decision was made. In every decision period, participants performed 4 blocks of 5 choices each. In each block, participants chose between the five snacks of the current run and a fixed money amount (1, 2, 3, or 4 CHF). Blocks were separated by a short instruction screen, indicating that the next 5 trials would offer the respective money amount. The order of the blocks and snack stimuli was randomized. The task was incentivized as participants were told in advance that a random trial would be selected after the experiment and they would receive their choice in that trial, be it a money amount or the snack. If no choice was made for this trial, they would not receive any option.

During the recall period we tested whether participants could recall the identicon-snack associations correctly. In each recall trial, we presented a red fixation dot in a white central square (jittered minimum fixation time as during encoding), succeeded by an identicon image. Participants pressed either the f or j button on the keyboard to indicate whether they could recall the snack correctly. Button-option assignment was counter-balanced across participants. A screen followed asking “Which snack?”. Participants verbally reported which snack they believed to be associated with the identicon. The experimenter noted the answer and initiated the source recall procedure. A screen asking “Which side?” was presented. Participants responded via button press, if they thought the snack was presented on the left (botton f) or on the right (button j) hemifield during the encoding period. Each recall period consisted of 5 trials, one for every identicon-snack association. The order of stimuli was randomized.

Participants were familiarized with (session 1) or remembered of (session 2) the experimental procedures in the beginning of each session. In both sessions, they performed a practice run before data collection started and were encouraged to ask questions before and after the practice run. We collected 7 runs of the memory condition (≙ 140 memory trials) and 3 runs of the value condition (≙60 value trials). Runs contained either memory or value trials. The order of runs was pseudo-randomized so that the first runs (session 1: runs 1 through 4, session 2: runs 1 and 2) contained only memory trials. This was done to increase participants’ familiarity with the task affordances. For the following runs, the order was randomized. The composition of the 5 snacks in each run was pseudo-randomized so that each run contained a snack of each willingness to pay quintile, obtained from the snack ratings.

#### 4.2.4 Post-experiment survey

After finishing the remember-and-decide task, participants rated their subjective familiarity with each snack. As for the snack ratings, participants moved a slider on a continuous scale to indicate how familiar they were with each snack. Again, the slider had to be moved at least once. Each snack familiarity was rated once. The order of the snacks was randomized.

After the familiarity rating task, participants would receive their incentives and were allowed to eat them during the following interview. In the semi-structured interview, participants indicated if they had specific strategies and/or difficulties while solving the tasks, and, if so, verbalized their respective strategies and/or difficulties. The experimenter took notes, and, when needed, asked clarification questions.

### 4.3 Trial selection

Data analysis in this paper focuses on trials from the decision period of the experimental session 2. We selected trials which fulfilled three criteria before pursuing data analysis. First, we selected trials in which we could assume that choice and RT data depended on a terminated drift-diffusion process. Therefore, we selected only trials where participants made a decision after 200 ms (48 trials) and before the response deadline (21 trials), coherent with our previous work (Kraemer et al., 2021a). Second, we excluded trials with excessive EEG noise (102 trials, see 4.5 for details of EEG preprocessing). Third, we wanted to compare value trials with memory trials, in which participants had access to the information of snack identity (in contrast to other work from our group, which investigated trials in which snack identities are only retrieved partially; cf. Kraemer et al., 2022). Thus, we rejected memory trials where participants could not recall the snack identity correctly during the recall period (636 trials). In total, we analyzed 2299 value trials (per participant *M* = 58.95, *s*.*d*. = 1.93, range: [51, 60], 98.25% of all value trials) and 4712 memory trials (per participant *M* = 120.82, *s*.*d*. = 15.97, range: [62, 134], 86.62% of all memory trials).

### 4.4 Behavioral modeling

Our behavioral analysis served two purposes: First, we wanted to know whether there are behavioral differences between value and memory trials. Second, we examined the suitability of the DDM for this task by testing whether its qualitative predictions are coherent with participants’ behavior. The DDM predicts that a) choices should follow a binary choice curve which depends on value difference of the choice options, and b) RT should describe an inverted U shape as a function of value difference (e.g., Clithero, 2018), or put differently: RT should decrease with absolute value difference of choice options.

To address these questions, we formulated two mixed effects regression models, henceforth labeled the choice model and the response time model. Both models were fitted in a hierarchical Bayesian framework (Lee and Wagenmakers, 2013) and had the functional form:

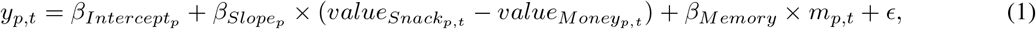

where *y*_*t*_ is the data (binary choice or log-transformed RT) of a trial *t. y*_*t*_ depended on participant-specific intercept and slope parameters 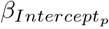 and 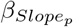, the difference of *z*-transformed subjective snack value and money amount, the fixed effect of trial type *β*_*Memory*_ which could affect the estimation in memory trials via the boolean variable *m*_*t*_ (*m* = 1, if *t* is a memory trial), and an unspecific error term *ϵ*. 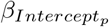 and 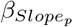 were participant-specific random effects which depended on group-level normal distributions 𝒩(*µ*_*Intercept*_, *σ*_*Intercept*_) and 𝒩(*µ*_*Slope*_, *σ*_*Slope*_). We specified standard-normal prior distributions for *µ*_*Intercept*_, *σ*_*Intercept*_, *µ*_*Slope*_, *σ*_*Slope*_ and *β*_*Memory*_. As variability parameters, *σ*_*Intercept*_, *σ*_*Slope*_ and *E* were constrained to be larger or equal zero. Importantly, the two models differed only with respect to a *logit* link function in the choice model, which mapped the model parameters to a binary choice outcome.

To estimate the models, we used a Markov-Chain Monte-Carlo (MCMC) approach for hierarchical Bayesian statistics using the probabilistic programming language *Stan* (Stan-Development-Team, 2018, *version 2*.*17*.*0). Four chains of 4,000 samples each were initialized of which 50% warm-up samples were discarded. The Gelman-Rubin* 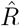 statistic (Gelman and Rubin, 1992) indicated convergence for all relevant parameters 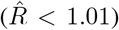. After having estimated posterior probability distributions of the model parameters, we performed statistical inference by calculating the 95% - Bayesian highest density intervals (Kruschke, 2015) and Bayes Factors. To calculate Bayes Factors, we applied the Savage-Dickey method to evaluate the inverse ratio of posterior and prior density at a null-effect level of zero (Lee and Wagenmakers, 2013). If this ratio is smaller than 1, evidence points toward the null-hypothesis that there is no effect. If it is larger than 1, evidence points toward the alternative hypothesis. The exact value of the Bayes factor specifies, how much more likely the alternative hypothesis is than the null hypothesis, given the model and the data.

### 4.5 EEG Preprocessing

We preprocessed EEG data with a custom-made script, drawing on the mne-Python library (Gramfort et al., 2013, version 0.17). We referenced the data to the average potential, obtained at the Mastoid electrodes. After applying a band-pass filter (between 1*/*7 Hz and 60 Hz) and a Notch-filter (at 50 Hz), we visually inspected trials for excessive noise. We fitted an independent components analysis and carefully selected components to exclude based on topography (spatially confined artifacts), periodicity (components that were only present in some blocks), and power spectrum (untypical drop in power spectrum). Further, components that were related to horizontal and vertical eye movements (as measured via EOG) were removed from the data.

We analyzed EEG data separately for stimulus-locked and response-locked epochs, covering the time windows [−700, 2000] ms relative to the snack/identicon presentation and [−1000, 100] ms relative to the participants’ response. We baseline-corrected all epochs to a time window of [−500, 100] ms relative to stimulus onset and down-sampled the data to obtain a temporal resolution of 8 ms. We further applied a current source density transformation to the data to improve topological localization of the signals.

### 4.6 EEG Analyses

To measure the temporal dynamics of the decision process, we focused on three properties of the LRP. The stimulus-locked LRP onset, the slope of the response-locked LRP and the response-locked LRP peak amplitude. To estimate these properties, we first calculated the empirical LRP using a double subtraction process, where the average effector-specific activity over the left and right motor cortices (electrodes C3 and C4) are subtracted (Coles, 1989).

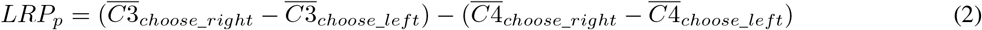

For every participant *p* and condition (memory and value trials), we calculated the LRP relative to the stimulus presentation and to the response. We applied a Gaussian smoothing kernel (s.d. = 24 ms) to the time-resolved LRP to deal with momentary signal noise.

We estimated the properties of the LRP with a segmented regression method (Schwarzenau et al., 1998) where the LRP time course follows a function *y*(*t*) = *f*(*t*) + *ϵ f*(*t*) describes the progression of the LRP as straight lines in a pre-onset and post-onset period.

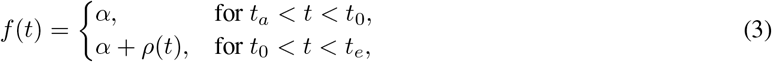

In the pre-onset period between the trial onset (*t*_*a*_) and the LRP onset *t*_0_, the signal is modeled as a horizontal line where the intercept *α* can be interpreted as pre-onset baseline activity. In the post-onset period between *t*_0_ and the LRP peak latency (*t*_*e*_), the signal is modeled as a straight line, originating at baseline level and progressing towards the peak amplitude with a slope *ρ*.

We fitted 2 × 2 (condition: memory and value; event-relation: stimulus-locked and response-locked) segmented regression models to the LRP time courses. *t*_*a*_ was 0 ms in stimulus-locked trials and -1,000 ms in response-locked trials. *t*_*e*_ was the peak latency of the respective LRP time course. We estimated the models in a hierarchical Bayesian framework, where *α*_*p*_, *ρ*_*p*_ and 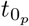 were random effects on the participant level which depended on group level normal distributions 𝒩(*µ*_*α*_, *σ*_*α*_), 𝒩(*µ*_*ρ*_, *σ*_*ρ*_) and 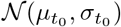. We specified standard-normal prior distributions for all population parameters. The variability parameters *σ*_*α*_, *σ*_*ρ*_ and 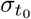 were constrained to be non-negative. The onset parameter 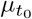 was constrained to be equal or larger than *t*_*a*_. We estimated the model in *Stan*, initializing four chains of 4,000 samples each (50% warm-up). 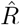 were smaller than 1.05 for all group-level parameters of the models, which indicated acceptable convergence of the chains.

To compare stimulus-locked LRP onsets and response-locked slopes, we computed posterior distributions of difference between memory and value condition. Posterior samples of stimulus-locked 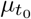 and response-locked *µ*_*ρ*_ in the value condition were subtracted from the corresponding samples in the memory condition. The ratio of samples larger than 0 can be interpreted as the probability, that the respective parameters were larger in the memory condition than in the value condition, given the model and the data. To compare response-locked peak amplitudes – which were not estimated by the segmented regression models – we performed a paired t-test on the measured minimum amplitudes within [−250, −50] ms relative to participants’ responses. We computed the Bayes Factors with the Python-based Pingouin library (Vallat, 2018, version 0.3.4) and specified the default Cauchy-prior with a scale parameter of 0.707.

We further sought to inform a neuro-cognitive model with individual estimates of the LRP properties onset, slope and peak (see 4.8). To deal with the relatively poor signal-to-noise ratio of EEG data on the participant level, we used a jackknifing method to obtain individual estimates of the LRP properties (Stahl and Gibbons, 2004). In a leave-one-participant-out procedure, we calculated the average LRP waveform and fitted a non-hierarchical Bayesian segmented-regression model to average LRP waveform (4 chains of 1,000 samples each; 50% warm-up; 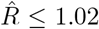 indicating good convergence). We did this for every participant, every condition and every event-relation. We used the resulting posterior mean estimates of stimulus-locked *t*_0_ as 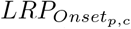, the response-locked posterior mean *ρ* estimates as 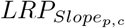 and the response-locked peak amplitudes of the waveforms as 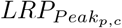. These estimates indicated how each individual contributed to the grand average LRP waveforms and thereby yielded estimates of individual variation for the participant. We *z*-standardized these estimates and embedded them as covariates in the neuro-cognitive model.

### 4.7 Cognitive Modeling

To investigate whether and how decision processes differ when information about value is sampled from memory, we applied the DDM (Ratcliff, 1978; Ratcliff and McKoon, 2008), arguably the most widely used sequential sampling model Ratcliff2016. Our drift diffusion model estimated the four parameters non-decision time *T*_*er*_, boundary separation *a*, starting point bias *z* and drift-rate *ν*. Choices and RT were thus assumed to follow a first-passage Wiener distribution.

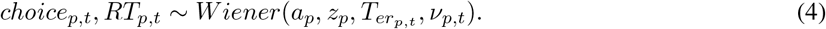

Whereas the parameters *a*_*p*_ and *z*_*p*_ were estimated for each individual (index p for participant), the parameters 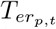 and *υ*_*p,t*_ additionally varied from trial to trial (index t for trial). In general, the diffusion parameters depended on a combination of random intercepts *β* which depended on group-level normal distributions 𝒩(*µ, σ*) (see table 1 for respective prior distributions). Critically, we equipped the model with additional fixed effects parameters (*δ* parameters) which modeled the difference between value and memory condition. A credibly positive (negative) *δ* can be interpreted as an increased (decreased) value of the corresponding diffusion parameter in the memory condition. At the same time, evidence for the null hypothesis of no condition difference can be tested by assessing the degree to which *δ* overlaps with zero. The boundary separation was modeled on the participant level as

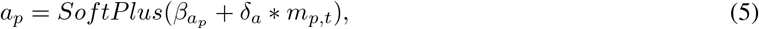

so that it depended on a random intercept 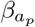 which was drawn from a group-level normal distribution 𝒩(*µ*_*a*_, *σ*_*a*_). *δ*_*a*_ was a fixed effect of the memory condition which affected the estimation during memory trials as indicated by the boolean dummy-variable *m*_*p,t*_. The sum of these terms was *SoftPlus*-transformed (*f* (*x*) = *ln*(1 + *e*^*x*^)), which, like the *exponential transformation*, constrains values to be positive, but has the advantage of ensuring a rather linear increase for higher parameter values. Starting point bias was estimated as a random effect where *z*_*p*_ = Φ(*β*_*z*_). The random intercept *β*_*z*_ was Phi-transformed to warrant that *z*_*p*_ is limited to values between zero and one. Non-decision time was estimated on the trial level so that the estimate was drawn from a normal distribution 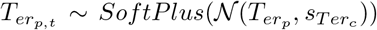. The center of the distribution was the participant-specific non-decision time 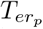, and the scale was the condition-specific non-decision time variability 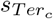 which was estimated for value and memory trials separately. We applied a SoftPlus-transformation to constrain the estimate to be positive. The participant-specific non-decision times was modeled as

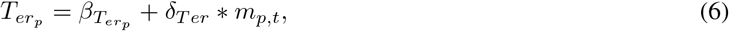

where 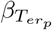 was a random intercept and *δ*_*Ter*_ was a fixed effect which specified the *T*_*er*_ difference in memory trials. Finally, the drift-rate was modeled as a scaled value difference of trial-specific snack- and money values (*z*-transformed) with an additional drift-rate bias term.

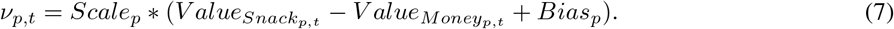

The drift-rate bias modeled the general preference of participants for either snack or money options, which was not explained with a starting point bias. We specified the bias as *Bias*_*p*_ = *β*_*Bias*_ + *δ*_*Bias*_, where *β*_*Bias*_ was a random intercept and *δ*_*Bias*_ was specified condition-specific difference of bias. The scale parameter *Scale*_*p*_ was modeled as

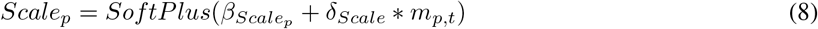

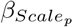 was a random intercept and *δ*_*Scale*_ was a fixed effect, specifying the condition-specific effect. We *SoftPlus*-transformed the sum to obtain strictly positive slope values.

The model was fitted in a hierarchical Bayesian framework in Stan (four chains of 4,000 samples each; 50% warm-up; 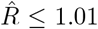). For statistical inference, we considered the 95%-HDI and Bayes Factors, which we computed with the Savage-Dickey method for *δ* parameters.

### 4.8 Neuro-cognitive Modeling

To investigate whether the LRP properties were related to the parameters of the DDM, we extended this model. We followed a *direct input* approach, where neuronal data was allowed to inform the parameters of the cognitive model (Turner et al., 2017). Individual LRP properties (estimated in the jackknifing procedure, see 4.6) were treated as covariates on the participant level which could explain variance via fixed effects *θ* parameters. A credibly negative *θ* parameter value implies – somewhat counter-intuitively – a positive correlation of individual LRP property and participant level DDM parameter value. The reason for this lies in the jackknifing procedure which omits the respective participant to estimate his/her effect on the grand-average (Stahl and Gibbons, 2004; Gluth and Meiran, 2019). If a *θ* parameter is indifferent from zero, one can infer that there is no relation of individual LRP properties and participant-level DDM parameter values.

The model was largely equivalent to the purely cognitive model (see 4.7). However, the parameters *T*_*er*_, *a* and *υ* were estimated with additional fixed effects.

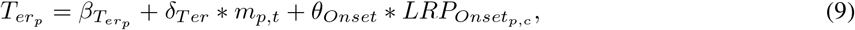

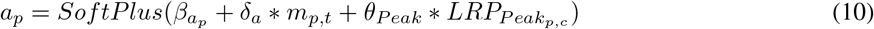

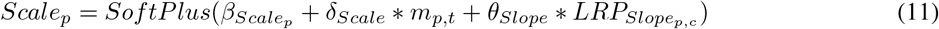

Thus, *θ*_*Onset*_ was a fixed effect which modeled the effect of the LRP onset in the respective condition. *a*_*p*_ de-pended on *θ*_*Peak*_ which covered the effect of LRP peak amplitude on boundary separation. Finally, *θ*_*Slope*_ was a fixed effect which modeled the effect of the response-locked LRP slope in the respective condition. All covariates 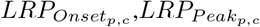 and 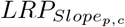 stemmed from the jackknifing procedure (see 4.6) and was calculated for each participant and condition (memory and value).

The model was fitted in a hierarchical Bayesian framework in Stan (four chains of 4,000 samples each; 50% warm-up; 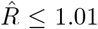). For statistical inference, we considered the 95%-HDI and Bayes Factors, which we computed with the Savage-Dickey method for *δ* and *θ* parameters.

## 5 Author Contributions

P.M.K and S.G. conceptualized the study design and methodology and administrated the project. P.M.K performed the software programming (experiment and analyses), computational and statistical analyses, conducted the experiments, curated the data, visualized the results and drafted the manuscript. S.G. validated the results, provided the resources, reviewed and edited the draft, supervised the work and acquired funding.

## 6 Acknowledgements

We thank Marie Habermann, Gregory Elbel and Noelle Burri for their support in data collection and Maria Wimber for helpful comments on data analysis. This work was supported by the Swiss National Science Foundation (SNSF Grant No. 172761 to Sebastian Gluth) and the Research Fund of the University of Basel (Grant No. 3PE1049 to Peter Kraemer).

## 8 Data and code availability

Data and code are uploaded on the Open Science Framework (https://osf.io/enwtd/) and are freely available.

## 9 Appendices

### 9.1 Appendix 1 - Identification of a memory signal

We conceptualized the remember-and-decide task not only to measure the LRP as a process of action selection, but also to identify a neural signature of episodic memory retrieval, unrelated to the decision process. To do this, we aimed to exploit the bihemispheric organization of the visual memory system (Gratton et al., 1997). When participants encode information in one visual hemifield, the information is represented by neural activity in early visual brain areas of the contralateral hemisphere. If the laterally encoded stimulus is retrieved from episodic memory, a reactivation process can be measured from the same brain areas which originally encoded the stimulus (Waldhauser et al., 2012, 2016). We reasoned that, if this procedure worked in the remember-and-decide task, we would be able to identify a neural signature of memory retrieval and study the temporal inter-relation of memory and decision processes.

During the encoding period, the identicon was presented in the center and the snack was presented either in the left or right visual hemifield. Participants maintained fixation on a red fixation dot in the center of the screen as ensured via online-eye tracking. When the identicon is presented in the decision period, we expected to see a signal of reactivation in those electrodes, which encoded the snack during the encoding period. To test this, we first identified electrode clusters which encoded visual information during the encoding period. In a second step, we tested if, during the decision period, those electrode clusters exhibited a hemisphere-specific reactivation, temporally locked to the presentation of the identicon.

For step one, we focused on the encoding period. We extracted epochs between -700 to 2.200 ms, relative to the stimulus onset. Following the procedures by Waldhauser et al. (2016), we selected electrodes posterior to the central channels (Fig. 6, grey) and applied a current source density transformation. For every participant and electrode, we computed the time frequency representation (TFR) of the signal between 1 and 30 Hz using Morlet wavelets. The TFR was truncated to a time window [−.5, 2]s and baseline-corrected to a time window [−.5, 0]s (both relative to stimulus onset). After applying a *z*-transformation, for every participant, we averaged the TFR of the trials where snacks were presented on the right and left hemifield and subtracted the values to obtain the hemifield-specific power difference Δ_*TFR*_ = *TFR*_*Snack*:*Left*_ − *TFR*_*Snack*:*Right*_. A positive value means that the electrode exhibits stronger desynchronisation when the snack was presented in the right hemifield vs. when it was presented in the left hemifield (and vice versa for negative values). Accordingly, electrodes in the left hemisphere represent visual information in the right hemifield with positive Δ_*TFR*_, and electrodes in the right hemisphere represent information with negative values. We tested, whether the electrodes encode the visual information by performing permutation cluster tests (1,000 permutations, critical *p* = .05). Those electrodes which contained a significant cluster were treated as encoding visual information. We identified two clusters which encoded visual information (Fig. 6, green).

**Figure 6:**
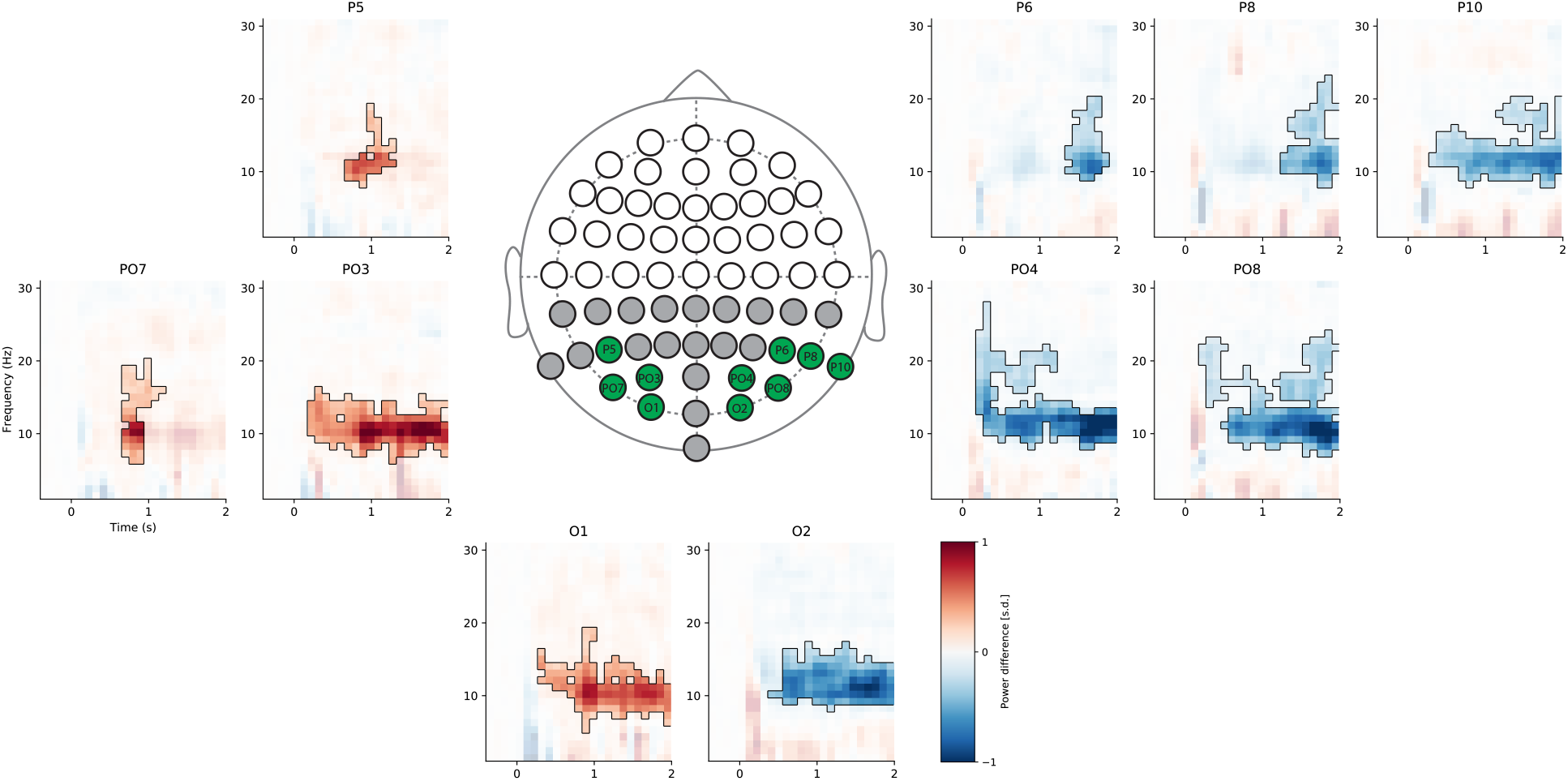
Identification of encoding electrodes. Plots show average power differences in time frequency representation (Δ_*TFR*_) between trials where snacks were encoded on the left vs. right hemifield. The drawing in the center pictures the electrodes. Grey electrodes depict the electrodes which were tested for significant Δ_*TFR*_ clusters. Green electrodes were found significant in permutation cluster tests.

Next, we tested, whether we see hemifield-specific reactivation in the found electrode clusters. We extracted epochs from the decision period ([−.7, 2.2]s; baseline corrected [−.5, .−1], both relative to stimulus onset; *z*-transformed). We averaged the power values for both clusters and computed Δ_*TFR*_ as above. Permutation cluster tests (1,000 permutations) indicated no hemisphere-specific reactivation during decision trials (best left cluster: *p* = .17; best right cluster: *p* = .8, see Fig. 7).

**Figure 7:**
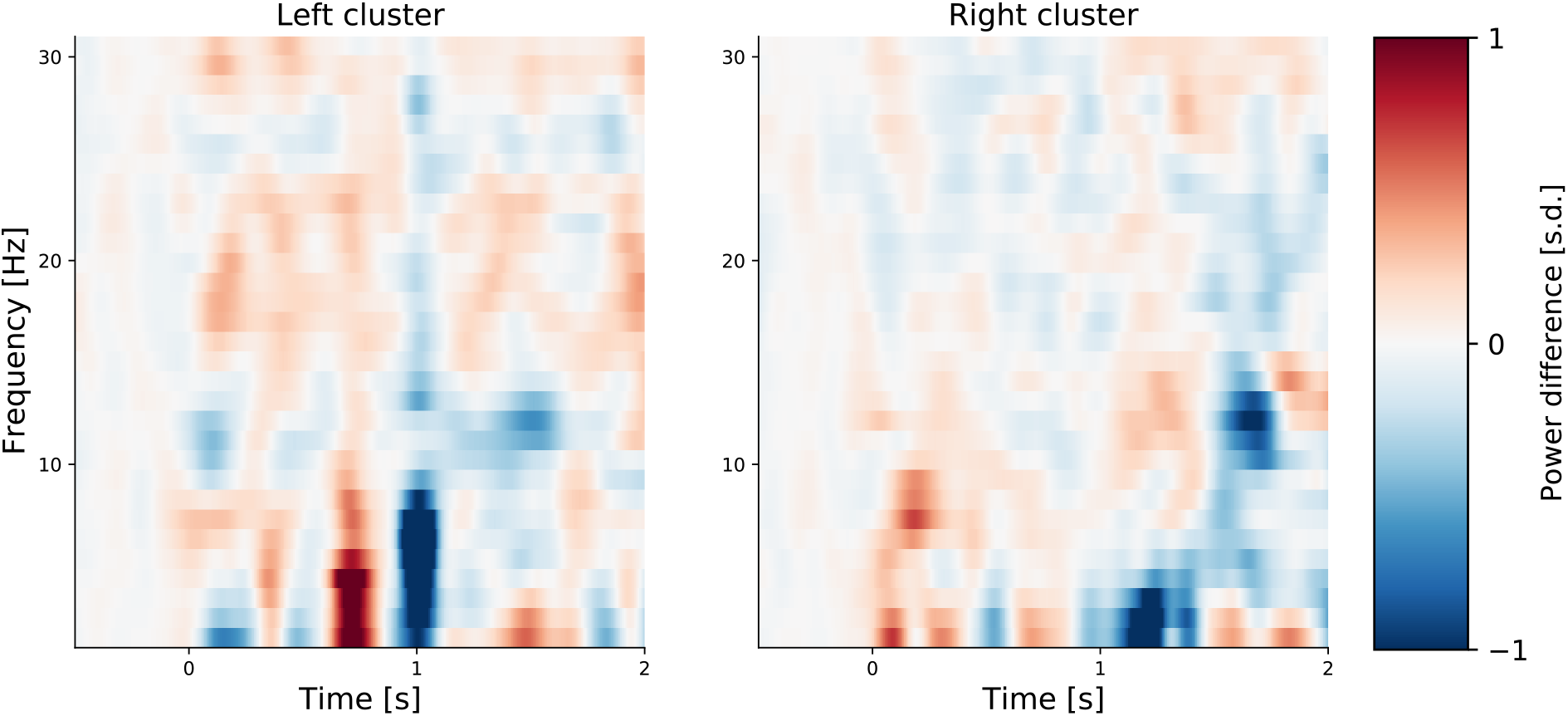
Test for reactivation in both clusters. Plot shows average power differences in time frequency representation (Δ_*TFR*_) in the decision period. Both, the left cluster (electrodes O1, PO3, PO7 and P5) and the right cluster (electrodes O2, PO4, PO8, P6, P8 and P10) show no statistically significant differences between trials, where the snack was encoded in the left vs. in the right visual hemifield.

In sum, whereas we were able to identify electrodes which encoded the location of the snack, these clusters did not exhibit hemifield-specific reactivation as we predicted, based on the literature (Waldhauser et al., 2012, 2016; Ede et al., 2019). An additional decoding exercise (comparable to Kerrén et al., 2018), where we aimed to decode hemifield-specific activity from trials in the decision period, did not indicate periods of reactivation. We assume that this null finding is related to different task demands between the reactivation paradigms in the literature and the remember-and-decide task. Whereas the stimulus location is relevant to solve the task in the studies Waldhauser et al. (2012, 2016); Ede et al. (2019), in the remember-and-decide task, participants only need to extract information about the snack identity, not in which visual hemifield it was presented. Future studies aiming to study the inter-relation of memory retrieval and decision making might adopt experimental designs that facilitate the identification of neural activity related to stimulus identity.

### 9.2 Appendix 2 - Centro-parietal positivity

In addition to the LRP component, we tested whether another EEG component might yield a relationship to decision-making mechanisms. Specifically, we looked at the centro-parietal positivity (CPP, O’Connell et al., 2012; Kelly and O’Connell, 2013), a component recorded over centro-parietal areas (O’Connell et al., 2012; Kelly and O’Connell, 2013), which has been proposed as a neural marker for built-up rate of decision variables (like the drift-rate in terms of the DDM) in perceptual decisions and is presumably related to the P300 component (Twomey et al., 2015). This idea was extended to consumer choices decisions (Pisauro et al., 2017), according to which there should be an build-up several hundreds of milliseconds relative to the time of response. To check, whether this was the case in our data – and therefore whether the CPP might yield information about evidence accumulation in the remember-and-decide task – we extracted EEG epochs ([−1, .1] relative to the response) of the CPz electrode and averaged them over trials and participants. As can be inspected in Figure 8, the CPP characteristic of an response-locked signal build-up was not present in our data. Also, another approach including the simulation of response-locked evidence accumulation traces and correlation with EEG activity [similar to]Pisauro2017,Polania2014 did not allow the identification of a reliable build-up. Realizing that we had, in comparison to Pisauro et al. (2017), only relatively few trials and comparatively variable RT (in our case, RT could be up to 7 s; Pisauro et al., 2017, had a time limit of 1.25 s), it is possible that we lacked statistical power to draw conclusions from this null-finding. We therefore refrained from further analyses, noting, however, that several recent studies suggest a relatively high task-specificity of the CPP (Lui et al., 2021; Frömer et al., 2021).

**Figure 8:**
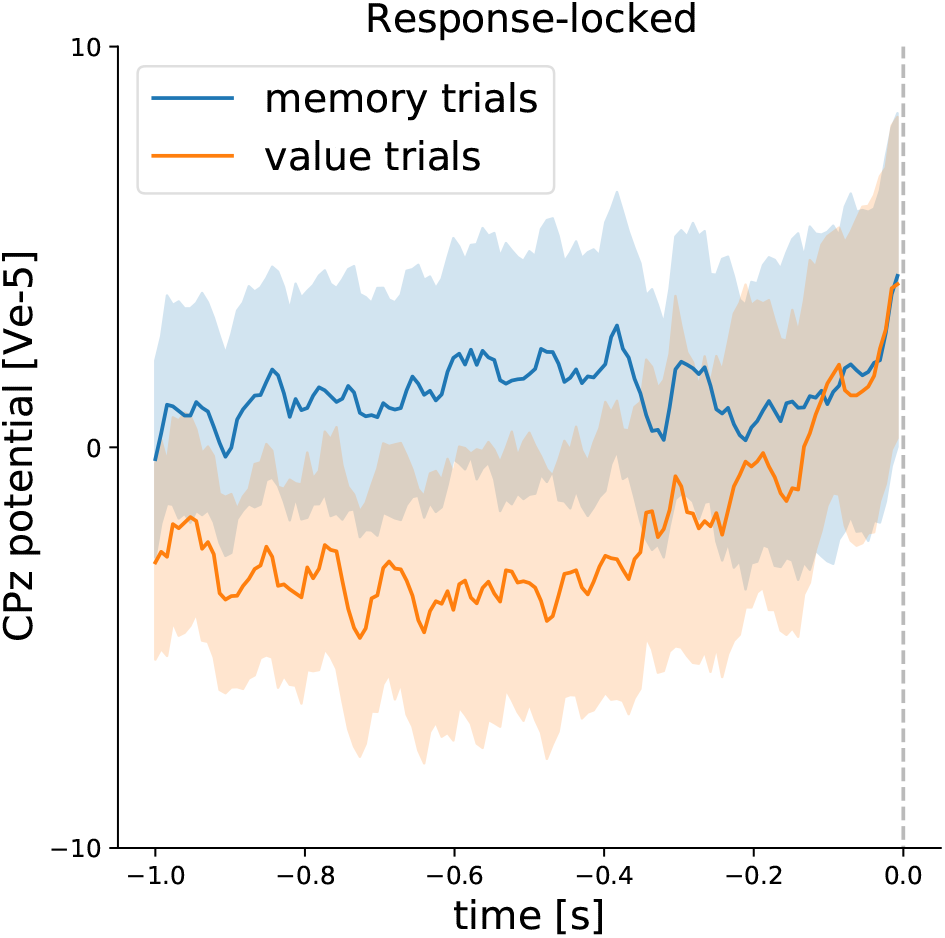
Mean Response-locked CPP signal for both conditions. Shaded areas indicate 95% bootstrapped confidence intervals. The vertical dashed line indicates the event of participant response.

